# Regeneration of Functional Retinal Ganglion Cells by Neuronal Identity Reprogramming

**DOI:** 10.1101/2020.07.16.203497

**Authors:** Xiaohu Wei, Zhenhao Zhang, Huan-huan Zeng, Xue-Feng Wang, Wenrong Zhan, Na Qiao, Zhen Chang, Lu Liu, Chengyu Fan, Ziwei Yang, Xiaoming Li, Yang Yang, Hongjun Liu

**Affiliations:** School of Life Science and Technology, ShanghaiTech University, Shanghai 201210, China; Institute of Biochemistry and Cell Biology, Shanghai Institutes for Biological Sciences, Chinese Academy of Sciences, Shanghai 200031, China; College of Life Sciences, University of Chinese Academy of Sciences, Beijing 100049, China; Institute of Neuroscience, Chinese Academy of Sciences, Shanghai 200031, China

**Keywords:** Retina, Retinal Ganglion Cell, Neuroregeneration, Neuronal Reprogramming, Regenerative Medicine, Glaucoma, Optic Neuropathy, Retinal Ganglion Cell Regeneration

## Abstract

Degeneration of retinal ganglion cells (RGCs) and their axons underlies vision loss in glaucoma and various optic neuropathies. There are currently no treatments available to restore lost vision in patients affected by these diseases. Regenerating RGCs and reconnecting the retina to the brain represent an ideal therapeutic strategy; however, mammals do not have a reservoir of retinal stem/progenitor cells poised to produce new neurons in adulthood. Here, we regenerated RGCs in adult mice by direct lineage reprogramming of retinal interneurons. We successfully converted amacrine and displaced amacrine interneurons into RGCs, and observed that regenerated RGCs projected axons into brain retinorecipient areas. They convey visual information to the brain in response to visual stimulation, and are able to transmit electrical signals to postsynaptic neurons, in both normal animals and in a diseased model. The generation of functional RGCs in adult mammals points to a therapeutic strategy for vision restoration in patients.

## INTRODUCTION

RGCs are the final output neurons of the retina that process visual information and transmit it to discrete brain visual areas to form vision (Sanes and Masland, 2015; Seabrook et al., 2017). Loss of RGCs is a leading cause of blindness in a group of diseases broadly categorized as optic neuropathies, including glaucoma, hereditary optic neuropathies, and disorders caused by toxins, nutritional defects and trauma (Quigley, 2016; Vetter et al., 2017). Vision loss in these patients is irreversible since humans and all mammals lack the ability to generate RGCs in adulthood (Jorstad et al., 2017). There is great interest in developing regenerative therapies to restore lost vision in these patients (Benowitz et al., 2017; Calkins et al., 2017; Crair and Mason, 2016; Laha et al., 2017; Marchetti et al., 2010; Roska and Sahel, 2018; Stern et al., 2018).

One attractive approach of developing regenerative therapies for optic neuropathies is to replace lost ganglion cells and reconnect the retina to the brain using endogenous cells (Crair and Mason, 2016; Goldberg et al., 2016; Vetter et al., 2017). Tremendous efforts have been made to identify retinal stem/progenitor cells and to understand how retinal neurons are generated in a variety of model organisms (Amato et al., 2004; Chiba, 2014; Dyer and Cepko, 2000; Fischer and Reh, 2000, 2001; Karl et al., 2008; Livesey and Cepko, 2001; Ooto et al., 2004; Tropepe et al., 2000; Yoshii et al., 2007). Previous studies demonstrated that lower vertebrates, like fish and amphibians, functionally regenerate their retinas following injury, and Müller glia are the cellular source of regenerated retinal neurons (Goldman, 2014; Hyde and Reh, 2014; Lamba et al., 2008). By contrast, Müller glia in mammals do not have this capacity and mammals, including humans, also do not have other reservoirs of retinal stem/progenitor cells poised to regenerate retinal neurons in the adult stage (Cicero et al., 2009; Langhe and Pearson, 2019). The current consensus is that there is normally little to no ongoing addition of neurons in the mature mammalian retina.

Here we sought to identify strategies that allow regeneration of RGCs using endogenous retinal cell sources. We show that other retinal neurons can be used as an endogenous cellular source for RGC regeneration. By ectopic expression of transcription factors essential for RGC differentiation, amacrine and displaced amacrine interneurons can be reprogrammed into RGCs in adult mice. Regenerated RGCs project axons into discrete subcortical brain regions. They response to visual stimulation and are able to transmit electrical signals into the brain, both under normal conditions and in an animal model of glaucoma, where original RGCs have been damaged by increased intraocular pressure.

## RESULTS

### Reprogram Lgr5^+^ Amacrine Interneurons into RGCs *in vivo*

We first used the *Lgr5^EGFP-IRES-CreERT2^; Rosa26-tdTomato* mouse strain to test whether RGCs could be regenerated from amacrine interneurons. Lgr5 is expressed in a subset of retinal amacrine interneurons located in the vitreous side of the inner nuclear layer (Figure 1A). When Lgr5^+^ amacrine interneurons are labeled with the tdTomato reporter and lineage traced in adult mice, very few tdTomato^+^ bipolar cells and horizontal cells could be detected a few months later (Figures 1B and 1C), consistent with previous observation that some Lgr5^+^ amacrine cells can turn off Lgr5 expression and, transdifferentiate into other retinal lineages (Chen et al., 2015), exhibiting limited regenerative potential. As mice age, a small number of Lgr5^+^ amacrine cells could be detected in the retinal ganglion cell layer, suggesting that they might be capable of migrating from the inner nuclear layer to the retinal ganglion layer (Figure 1D). Some of these cells turn off Lgr5 expression, but they never turn into RGCs.

**Figure 1.**
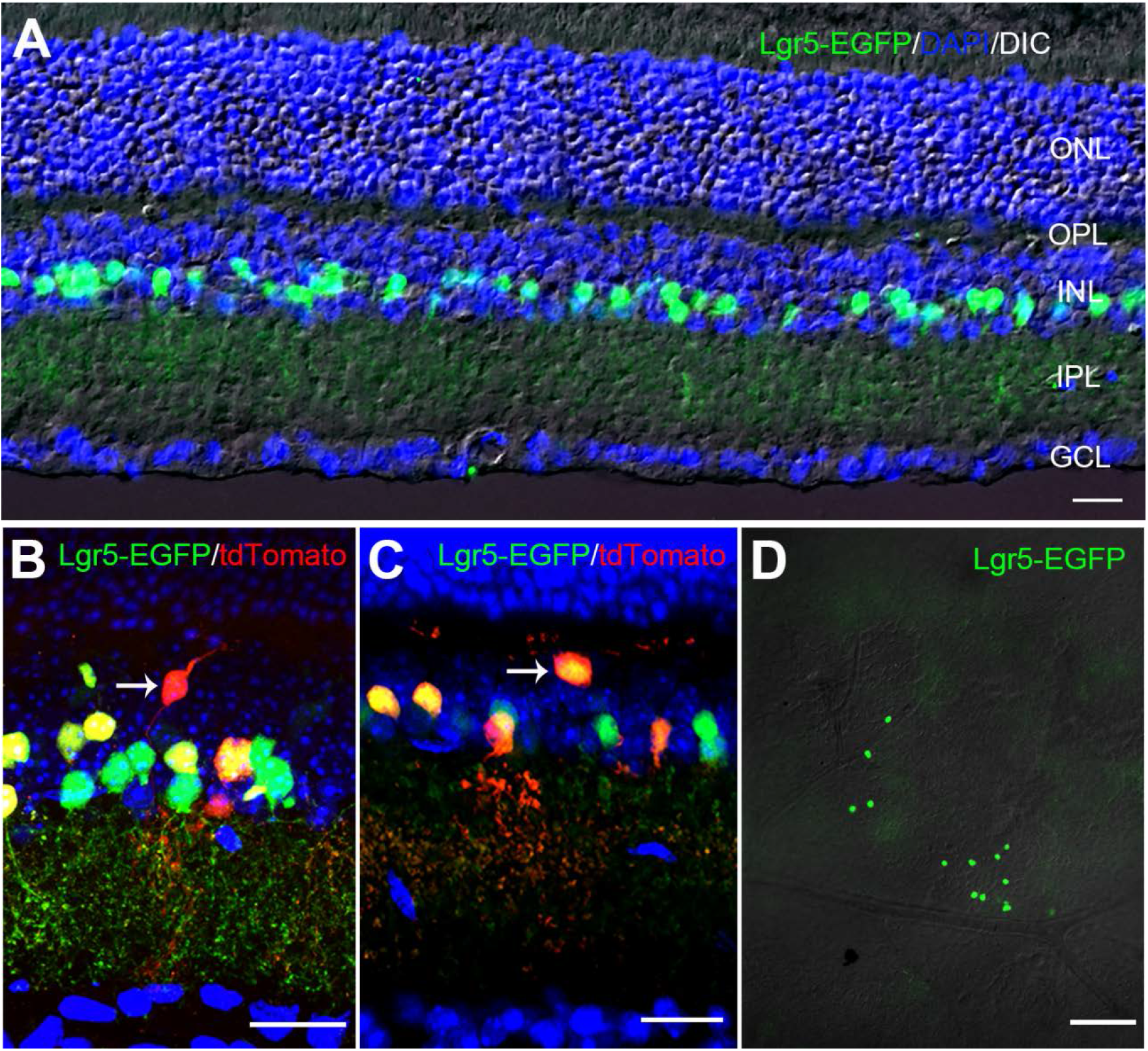
Lgr5^+^ Amacrine Interneurons Transdifferentiate into Other Neuronal Subtypes in Adult Mice. (A) Image of retina cross section showing Lgr5^+^ amacrine interneurons in the inner nuclear layer. (B and C) Images of retina cross sections from *Lgr5^EGFP-IRES-CreERT2^; Rosa26-tdTomato* mice. Arrows highlight generation of bipolar (B) and horizontal (C) cells from Lgr5^+^ amacrine interneurons. (D) Image of flat-mount retina sample, focusing on the retinal ganglion cell layer. ONL: outer nuclear layer; OPL: outer plexiform layer; INL: inner nuclear layer; IPL: inner plexiform layer; GCL: ganglion cell layer. Scale bar = 30 μm in panels A, B, and C. Scale bar = 200 μm in panel D.

To investigate whether Lgr5^+^ amacrine interneurons could be reprogrammed into RGCs, we devised an *in vivo* lineage tracing and reprogramming strategy (Figures S1A and S1B). We first labeled Lgr5^+^ amacrine neurons with the Rosa26-tdTomato reporter, and then ectopically expressed genes essential for RGC fate determination specifically in these cells (Badea et al., 2009; Brown et al., 2001; Gan et al., 1996; Jiang et al., 2013; Liu et al., 2001; Wu et al., 2015), using the Cre-dependent double-floxed inverted open reading frame (DIO) expression system delivered via adenovirus-associated virus (AAV). We analyzed the generation of RGCs from Lgr5^+^ amacrine cells by examining the presence of tdTomato^+^ cells with RGC morphology in flat-mount retina samples, and the presence of tdTomato^+^ axons in optic nerves at later time points. We did not observe any tdTomato^+^ RGC cells in flat-mount retina samples and tdTomato^+^ axons in optic nerves from control mice intravitreally injected with AAV-DIO-EGFP (Figures S1C and S1D). However, tdTomato^+^ cells with RGC morphology could be detected in retina samples from mice injected with AAVs, expressing a set of genes essential for RGC fate determination (Atoh7, Brn3B, Sox4, Sox11 and Isl1). Six weeks after induced gene expression, tdTomato^+^ cells with RGC morphology were observed in retina samples (Figures 2A and 2B, and Figures S1E-S1G). These cells projected axon-like projections towards the optic disc and extended into the optic nerve (Figure 2B and Figures S1E and S1F). Their cell bodies were located in the retinal ganglion cell layer, and can be stained with RGC-specific markers RBPMS and Brn3A (Figures 2C and 2D). Therefore, these cells could be potentially considered as newly generated RGCs.

**Figure 2.**
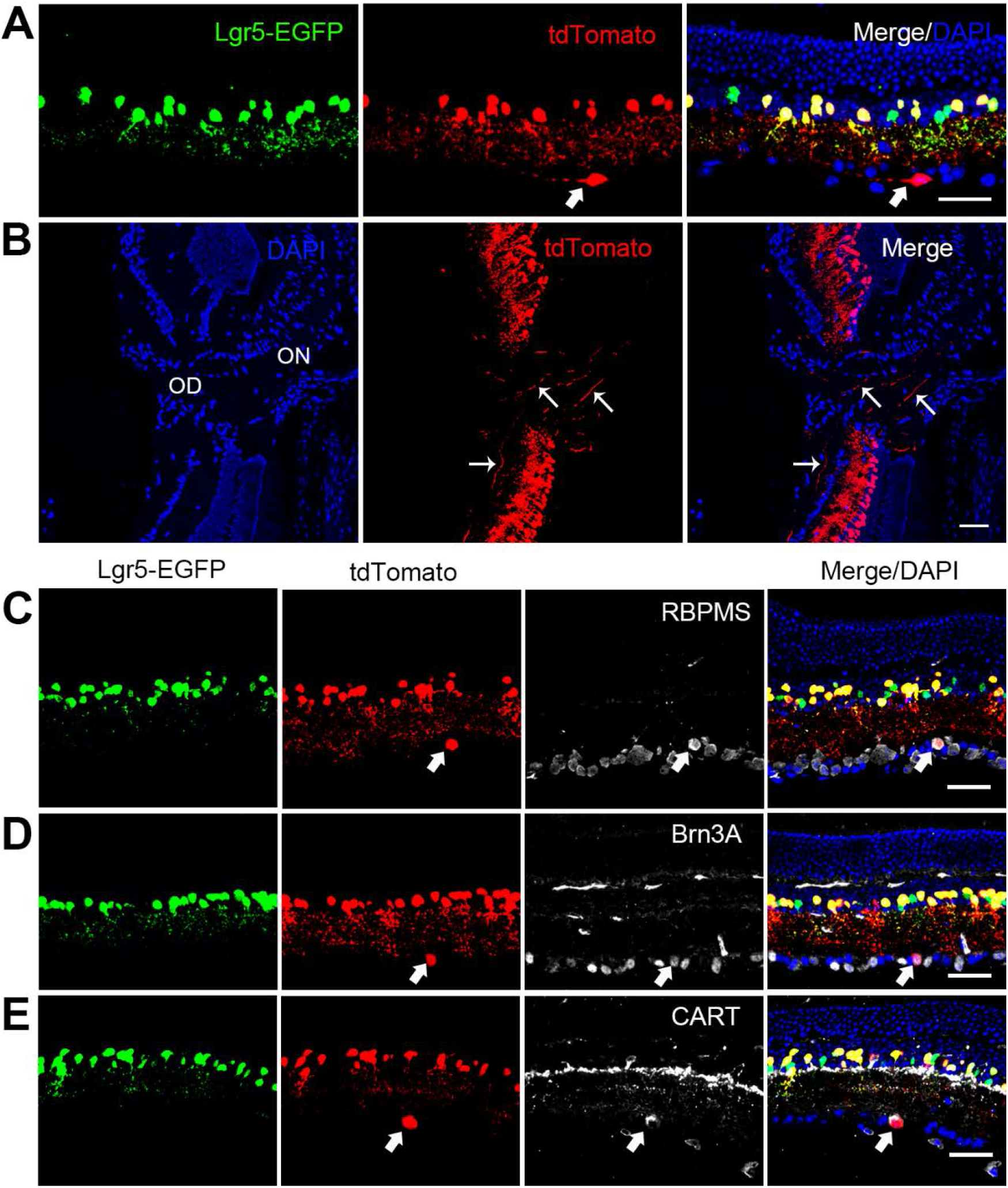
Reprogram Lgr5^+^ Amacrine Interneurons into RGCs *in vivo*. (A) Confocal image of retina cross section. Arrows highlight a regenerated RGC with an axon-like process. (B) Regenerated RGCs extend axons into the optic disc (OD) and optic nerve (ON). (C-E) Immuno-histological stainings of regenerated RGCs with antibodies specific for RBPMS (C), Brn3A (D) and CART (E). Scale bar = 40 μm.

On average, about 180 new RGCs per retina are regenerated 6 weeks after viral injection. This number is much higher than that of Lgr5^+^ amacrine cells present in the retinal ganglion layer when *in vivo* reprogramming was initiated (about 10-15 cells in the retinal ganglion cell layer of 2-3 month-old mice). This result suggests that ectopic expression of RGC fate-determining factors in Lgr5^+^ amacrine cells might trigger the migration of some of these cells from the inner nuclear layer to the ganglion cell layer. In support of this notion, tdTomato^+^ cells with lower Lgr5-EGFP expression level were detected in the inner plexiform layer (Figures S2A-S2C).

We tested the reprogramming activities of single transcription factors and their combinations and found that, even single transcription factor (Brn3B or Sox4) was capable of reprogramming Lgr5^+^ amacrine interneurons into RGCs, but with very low efficiency (Figure S2H).

Combination of Brn3B and Sox4 dramatically synergized reprogramming activity. Addition of Atoh7 to the Brn3B+Sox4 combination did not further improve reprogramming efficiency much (Figure S2H). Therefore, we used the Brn3B+Sox4 combination for the rest of experiments unless otherwise noted.

RGCs are a heterogeneous type of retina neurons that can be classified into distinct subtypes. We performed immuno-histological analysis with subtype-specific antibodies and found that, regenerated RGCs could be identified with anti-CART (for ON OFF directionally selective ganglion cells) and anti-SMI-32 (for α ganglion cells) (Figure 2E, and Figures S2D-S2G), but we did not detect any melanopsin-expressing intrinsically photosensitive ganglion cells. Together, these results suggest that ectopic expression of specific transcription factors is capable of reprogramming Lgr5^+^ amacrine interneurons into RGCs, and regenerated RGCs are subtype-specific.

### Regenerated RGCs Project Axons into Visual Nuclei in the Brain

To determine whether regenerated RGCs could rewire appropriately in the brain, we examined the axons of regenerated RGCs along the retinofugal pathway and their projections to the main brain retinorecipient areas. Six weeks after viral injection, many axons of regenerated RGC have traversed the entire optic nerve, passed the optic chiasm, and navigated into visual nuclei in the brain, including the dorsal and ventral lateral geniculate nucleus (dLGN and vLGN), the pretectal area, and the superior colliculus (SC) (Figure 3). We did not observe aberrant projections of regenerated RGC axons to brain regions unrelated to the visual pathway.

**Figure 3.**
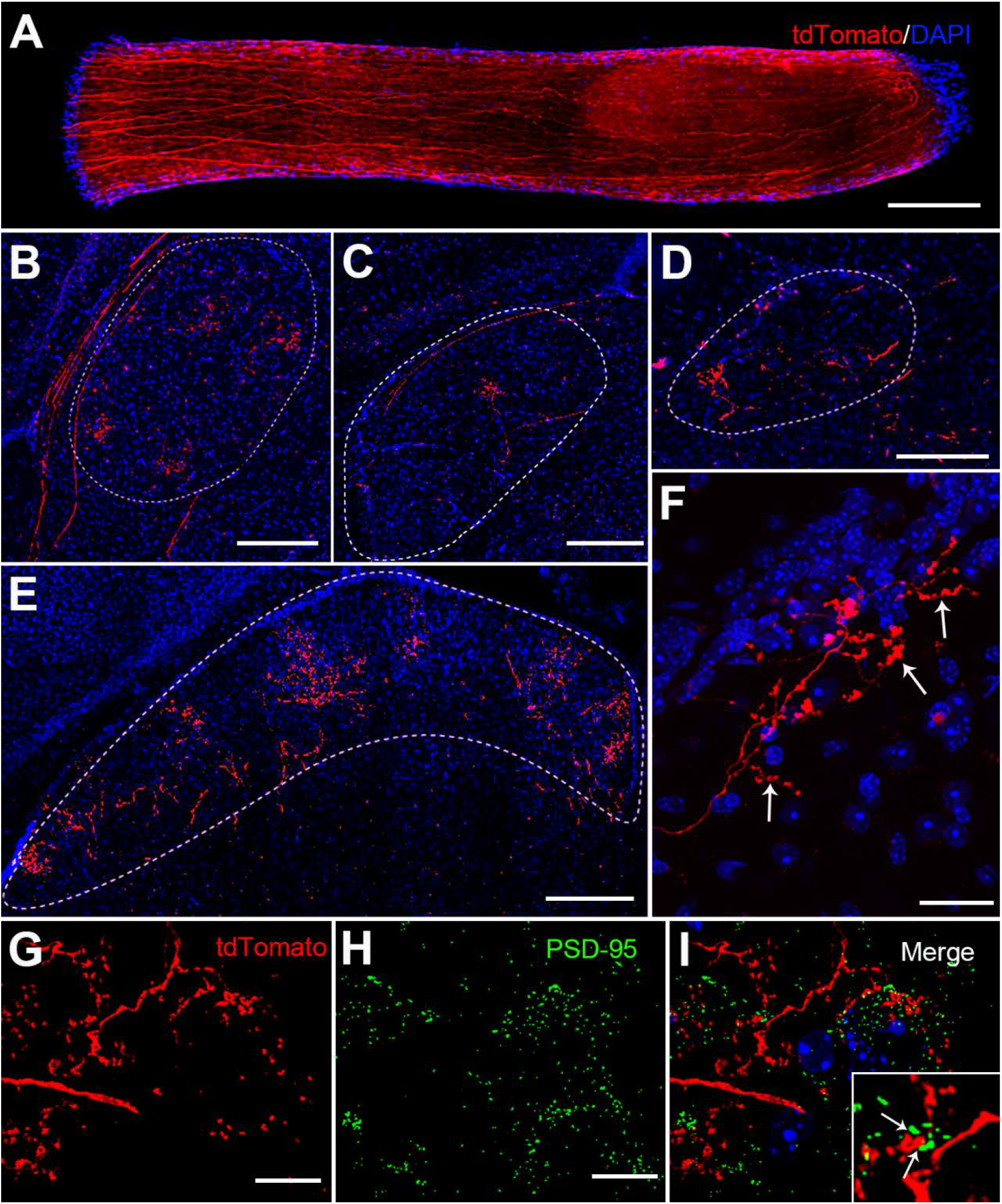
Regenerated RGCs Project Axons into the Brain. (A) Axons of regenerated RGCs within the optic nerve. (B-F) Projections of regenerated RGC axons in brain visual areas, showing tdTomato^+^ axon terminals in dorsal and ventral lateral geniculate nucleus (dLGN and vLGN, panels B and C respectively), olivary pretectal nucleus (OPN, panel D), and the superior colliculus (SC, panels E and F). Arrows in panel F highlight bouton-like structures on regenerated RGC axon terminals in the SC region. (G-I) PSD-95 staining of SC brain sections. Scale bars = 200 μm in panels A to E, = 40 μm in panel F, and = 10 μm in panels G and H.

Within retinorecipient areas, micron-sized varicosities are observed along axonal arborizations of regenerated RGCs. These varicosities are in close apposition to staining for the postsynaptic density protein PSD-95, suggesting that they are putative presynaptic boutons (Figures 3G-3I). Together, these data demonstrate that Lgr5^+^ amacrine interneuron-derived RGCs are capable of projecting axons into appropriate brain areas, establishing retina-brain connections.

### Reprogram Prokr2^+^ Displaced Amacrine Interneurons into RGCs

We think other retinal neurons could be reprogrammed into RGCs too. Displaced amacrine interneurons could serve as a better cellular source for RGC replacement, since they are located in the RGC layer. To test if this neuronal subtype could be reprogrammed into RGCs, we generated a *Prokr2^CreERT2^* knock-in mouse line (Figure S3A). The *Prokr2^CreERT2^* mice express the tamoxifen-inducible CreERT2 recombinase under the endogenous transcriptional control of the Prokr2 gene, which is expressed in a subgroup of displaced amacrine interneurons. As expected, in adult *Prokr2^CreERT2^; Rosa26-tdTomato* mice treated with tamoxifen, tdTomato^+^ cells are located in the retinal ganglion cell layer. They do not have optic projections and do not express the RGC maker RBPMS (Figures 4A and 4B, and Figures S3B-S3D).

**Figure 4.**
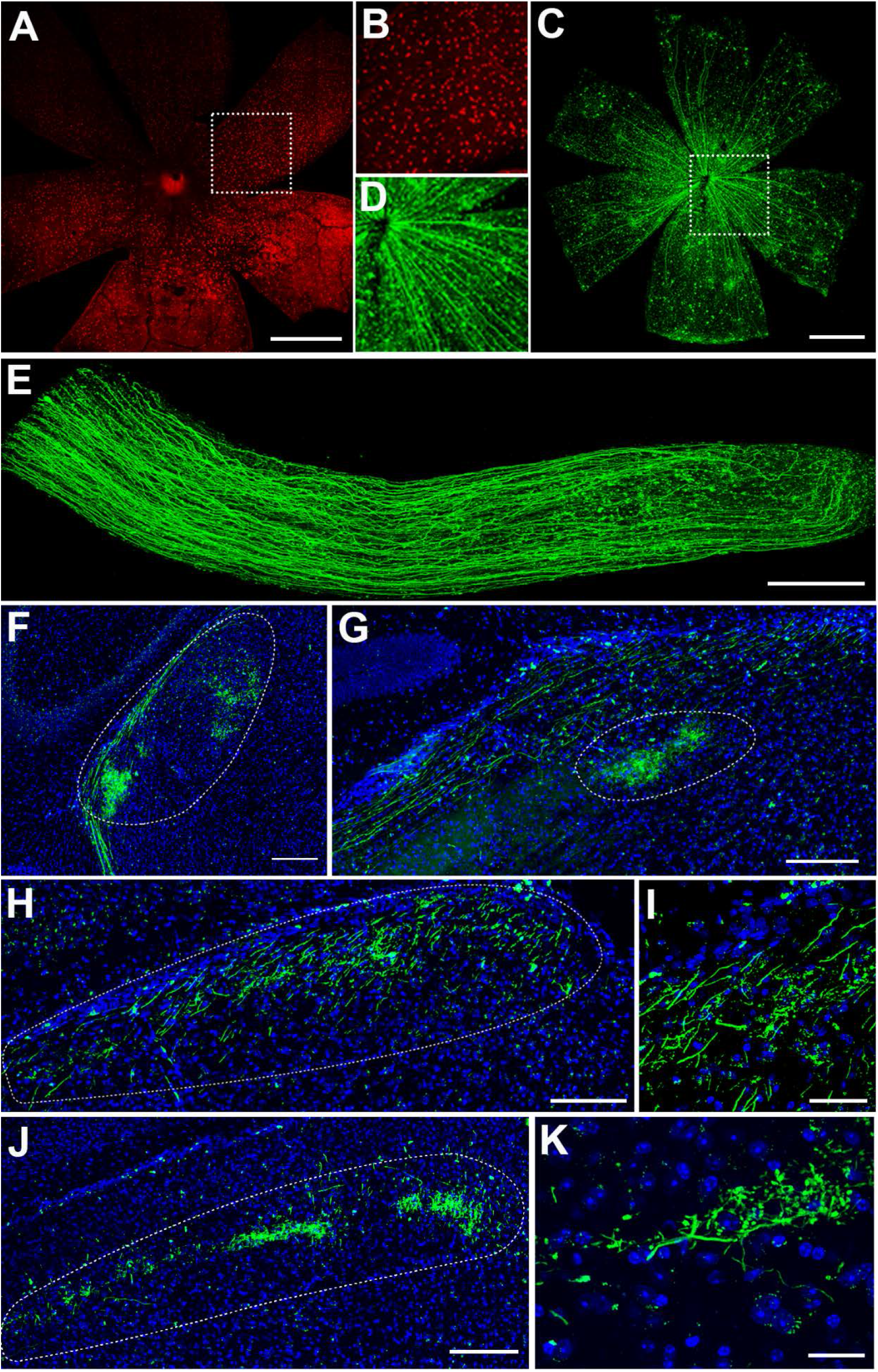
Reprogram Prokr2^+^ Displaced Amacrine Interneurons into RGCs. (A) Confocal image of flat-mount retina sample from *Prokr2^CreERT2^; Rosa26-tdTomato* mice. tdTomato^+^ displaced amacrine cells are shown in red. (B) Highlight of an area in panel A. (C) Confocal image of flat-mount retina sample from Prokr2^CreERT2^ mice injected with AAVs expressing transcription factors and EGFP. Regenerated RGCs extend axons to the optic disc. (D) Highlight of an area in panel C. (E) Axons of regenerated RGCs within optic nerve. (F-K) Projections of regenerated RGC axons to various brain retinorecipient areas, including dLGN and vLGN (F), OPN (G), and upper (panels H and I) and lower layer (panels J and K) of SC regions. Images in panels I and K are high-magnification views. Scale bars = 1000 μm in panels A and C, = 200 μm in panels E to G, and panel J, = 100 μm in panel H, and = 40 μm in panels I and K.

In addition to being expressed in the retina, Prokr2 is also expressed in cells of the optic nerve and the brain (Figures S3E-S3G). This prevented us from using the *Rosa26-tdTomato* reporter to track axons of regenerated RGCs. To overcome this obstacle, we labeled regenerated RGCs by co-expressing EGFP with transcription factors during programming (Figures S4A and S4B). We used two combinations of transcription factors (Brn3B+Sox4 and Atoh7+Brn3B+Sox4) for reprogramming and found that, both combinations could efficiently reprogram Prokr2^+^ displaced amacrine interneurons into RGCs. However, unlike in Lgr5^+^ amacrine interneurons, inclusion of Atoh7 to the Brn3a+Sox4 combination dramatically enhanced reprogramming efficiency (Figure S4C). Prokr2^+^ displaced amacrine interneuron-derived RGCs also extended axonal projections into the optic nerve and various brain visual targets (Figures 4E–4K). Thus, these results demonstrated that RGCs could be regenerated by reprogramming multiple retinal neuron subtypes *in vivo*.

### Regenerated RGCs Convey Visual Information to the Brain

To investigate whether regenerated RGCs could respond to visual stimulation and convey visual information to downstream targets in the brain, we labeled regenerated RGCs with the calcium indicator GCamp6f in *Lgr5^EGFP-IRES-CreERT2^* mice by adding AAV-DIO-GCamp6f to the reprogramming cocktail. We then exposed SC of anesthetized mice six weeks after viral injection, and used *in vivo* functional calcium imaging to measure the visually evoked calcium dynamics of regenerated RGC axon terminals.

When mice were presented with drifting gratings, individual RGC boutons along the axonal arborization in SC exhibited stimulus-evoked calcium signal, indicating that regenerated RGCs could respond to visual stimulation and transmit visual signals to the brain (Figure S5). Visually responsive boutons could be classified into distinct categories based on their response patterns. Boutons responded differently to on and off of stimulation, as well as to the orientation and direction of drifting gratings (Figures 5A–5D). Together, these data suggest that regenerated RGCs are capable of conveying visual information to the brain, and functionally distinct RGC subtypes could be generated by *in vivo* reprogramming.

**Figure 5.**
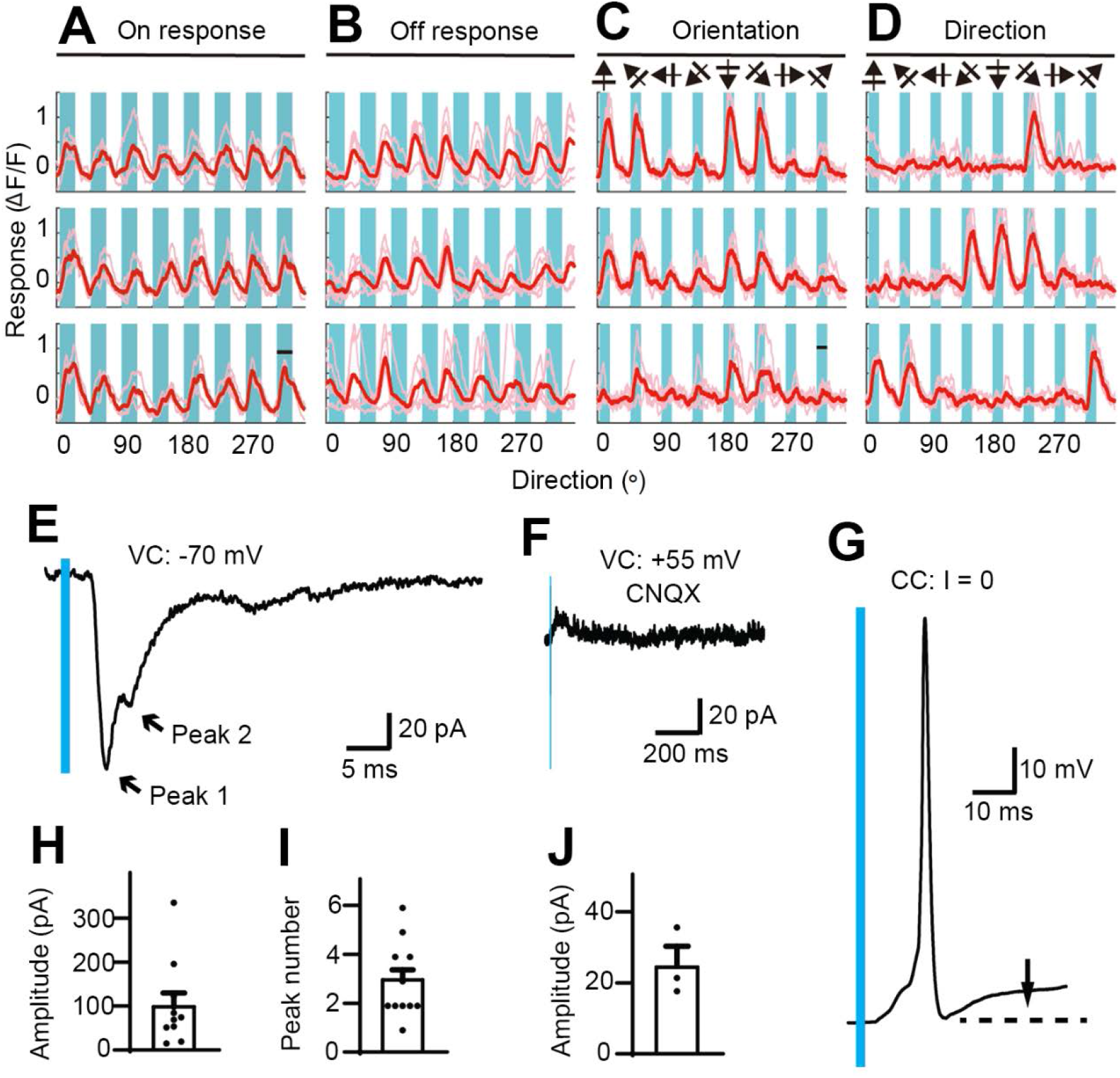
Regenerated RGCs Transmit Visual Information to the Brain and Establish Functional Connections with Postsynaptic Neurons. (A) Traces of calcium signals of three example axonal terminals in response to drifting gratings of different directions. Cyan patches mark periods of stimulus presentation, and the values on bottom indicate stimulus direction. Note that the three terminals shown in this panel have robust “on” responses. (B) Same plots as in panel A except that terminals shown here have robust “off” responses. (C and D) Traces of calcium signals of three axon terminals that have orientation selectivity (C) and direction selectivity (D). (E) An representative EPSC of light-evoked postsynaptic AMPA receptor responses in a SC neuron. Arrows indicate the postsynaptic response of multi-presynaptic inputs. (F) An representative EPSC of light-evoked postsynaptic NMDA receptor response in a SC neuron. (G) An example of light-evoked postsynaptic action potential in a SC neuron. (H and I) Summary of EPSC amplitudes (H) and peak numbers (I) of light-evoked AMPA receptor responses (n=11 from 7 animals). (J) Summary of EPSC amplitudes of light-evoked NMDA receptor responses (n=3 from 3 animals). Data are mean ± sme.

### Regenerated RGCs Establish Functional Synaptic Connections with Postsynaptic Neurons

To investigate whether regenerated RGCs could transmit neuronal signals to postsynaptic neurons in the brain, we expressed Channelrhodopsin-2 (ChR2) in regenerated RGCs in *Lgr5^EGFP-IRES-CreERT2^; Rosa26-tdTomato* mice, and used whole-cell patch recording to detect light-evoked postsynaptic responses of SC neurons on brain slices 8-10 weeks after viral injection.

When axon terminals of regenerated RGCs were stimulated with light, AMPA receptor-mediated excitatory postsynaptic currents (EPSCs) were detected in SC neurons. A single light impulse evoked AMPA receptor-mediated EPSCs with multiple peaks (Figures 5E, 5H, and 5I), suggesting that regenerated RGC axons formed multi-input synapses with SC neurons and activated AMPA glutamatergic receptors. NMDA receptor-mediated EPSCs and action potential were also detected in postsynaptic SC neurons after light stimulation (Figures 5F, 5G, and 5J). Together, these results suggest that, in response to light stimulation, regenerated RGC axon terminals release glutamate as neurotransmitter and establish functional synaptic connections with SC neurons.

### Regenerate Functional RGCs in a Mouse Model of Glaucoma

We next asked whether regenerated RGCs could repair visual circuits under diseased conditions. We are particularly interested in determining whether regenerated RGCs could still send axons to appropriate brain targets, and rebuild the retina-brain connection, when original RGCs and their axons have undergone degeneration.

We used an intraocular pressure increase-induced glaucoma model to damage RGCs and their axons, and optimized a condition that could cause degeneration of all RGC axons (Figures S6A and S6B). However, this damage condition also caused dramatic loss of Lgr5^+^ amacrine cells seven days after intraocular pressure increase (Figure 6B). Therefore, we searched for reagents that could protect Lgr5^+^ amacrine cells and found that, the Rock inhibitor Ripasudil could efficiently preserve Lgr5^+^ amacrine cells (Figure 6C).

**Figure 6.**
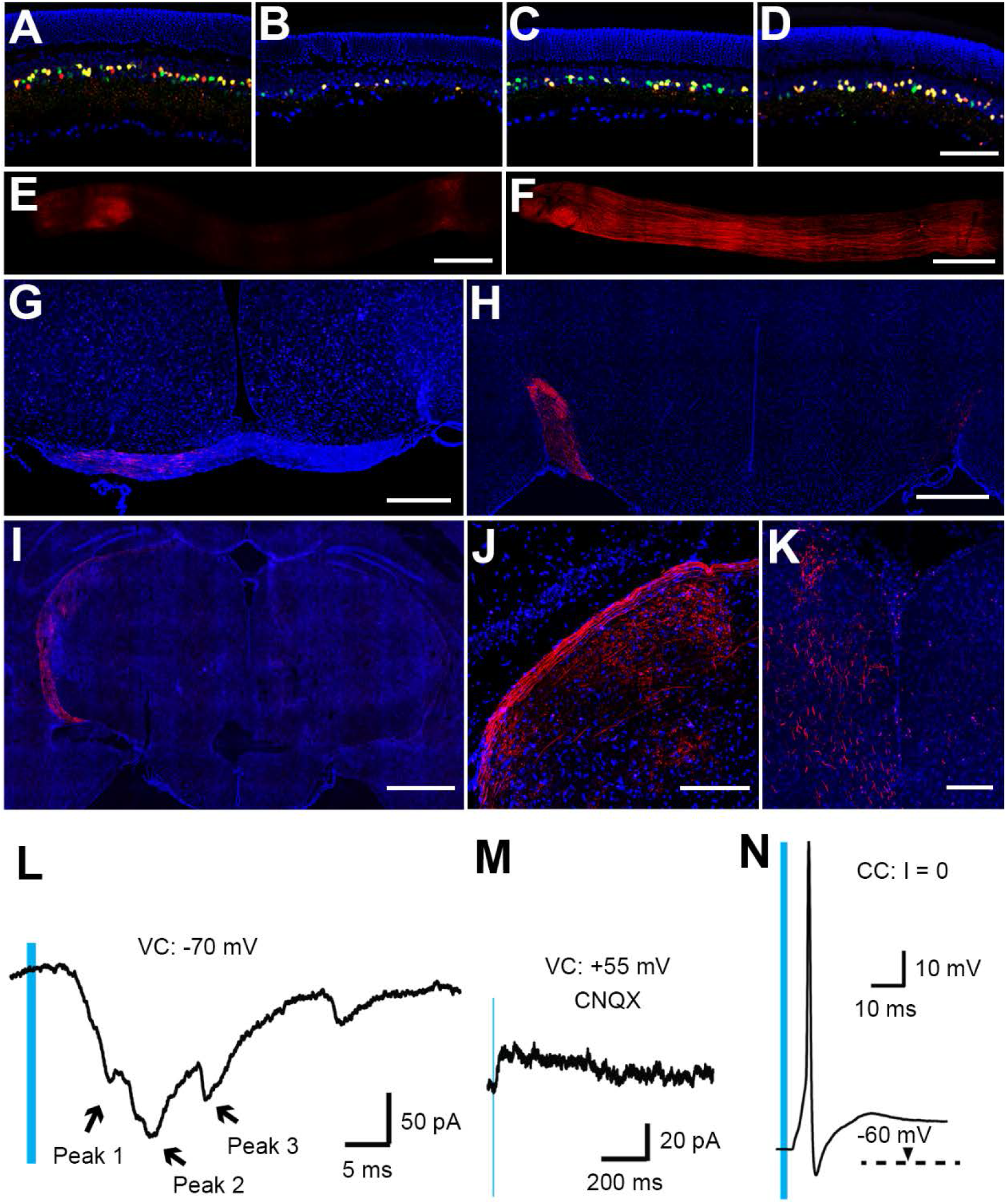
Regenerate Functional RGCs in a Mouse Model of Glaucoma. (A-D) Confocal images of retina samples. (A) Retina of normal *Lgr5^EGFP-IRES-CreERT2^; Rosa26-tdTomato* mice. (B) Retina of mice damaged by intraocular pressure increase (IPI) seven days ago. (C) Retina of mice damaged by IPI seven days ago but received daily Ripasudil treatment. (D) Retina of mice damaged by IPI but received both Ripasudil treatment and injection of AAVs expressing RGC fate-specification transcription factors (AAV-DIO-TFs). Mice were sacrificed 6 weeks after AAV injection. (E and F) Confocal images of optic nerves from eyes receiving AAV-DIO-EGFP (E) and AAV-DIO-TFs (F). (G-K) Confocal images of brain sections from *Lgr5^EGFP-IRES-CreERT2^; Rosa26-tdTomato* mice, which had received injections of AAV-DIO-EGFP in the left eye and AAV-DIO-TFs in the right eye. The majority of regenerated RGC axons were projected to the contralateral (left) side of the brain, with a small portion to the ipsilateral (right) side. Brain visual areas presented are optic track immediately after the optic chiasma (G), optic track (H), dLGN, vLGN and projection to the pretectal areas (I), dLGN (J), and SC (K). (L-N) Light-evoked postsynaptic responses of SC neurons, including AMPA receptor-mediated EPSC (L), NMDA receptor-mediated EPSC (M), and action potential (N). Scale bars = 100 μm in panels D, J and K, =200 μm in panels E and F, = 400 μm in panel G, and = 600 μm in panels H and I.

We devised a protocol that combined neuronal protection with *in vivo* reprogramming, to test if new generated RGCs could repair damaged visual circuitry in the *Lgr5^EGFP-IRES-CreERT2^; Rosa26-tdTomato* mice (Figure S6C). We damaged both eyes of *Lgr5^EGFP-IRES-CreERT2^; Rosa26-tdTomato* mice that have received tamoxifen injections to further label Lgr5^+^ amacrine cells with the tdTomato reporter. After damage, both eyes were treated with Ripasudil once a day, and seven days later, the left eye was injected with AAV-DIO-EGFP as control, whereas the right eye was injected with a combination of AAVs expressing transcription factors including Brn3B, Sox4 and Atoh7, to reprogram Lgr5^+^ amacrine interneurons. Six weeks after viral injection, mice were sacrificed for analysis. In the left eye injected with AAV-DIO-EGFP, no RGCs were regenerated, since no tdTomato^+^ RGC axons could be detected in the left optic nerve (Figure 6E). In contrast, overexpression of Brn3B, Sox4 and Atoh7 reprogrammed Lgr5^+^ amarine cells into RGCs, and regenerated RGCs projected tdTomato^+^ axons into the optic nerve and various brain visual targets of the contralateral side (Figures 6F-6K).

Regenerated RGCs established functional synaptic connections with postsynaptic brain neurons under diseased conditions. Light-evoked postsynaptic responses were detected in SC neurons on brain slices, where all RGC axon terminals were from regenerated RGCs after original ones had been damaged (Figures 6L-6N). Together, these results suggest that regenerated RGCs could reconnect the retina to the brain and transmit visual information to postsynaptic neurons even under diseased conditions.

## DISCUSSION

Our results demonstrate that functional RGCs can be generated in adult mammals by *in vivo* reprogramming of fully differentiated retinal interneurons. By ectopic expression of essential transcription factors, both amacrine and displaced amacrine interneurons can be precisely reprogrammed into RGCs, and new generated RGCs integrate into the visual circuitry and transmit visual information to the brain. Although *in vivo* neuronal identity reprogramming has been achieved in other regions of the central nervous system (CNS), successful conversion between neuronal subtypes was only restricted to the first postnatal week (De la Rossa et al., 2013; Rouaux and Arlotta, 2013), or with limited success in adult mice when the chemical compound valproic acid is present (Niu et al., 2018). In contrast, we demonstrate that, even without chemical-stimulant, retinal neuronal identity switching can be achieved in adulthood, and successful reprogramming even triggers migration of amacrine interneurons from the inner nuclear layer to the RGC layer. These results implicate that neurons exhibit surprisingly unexpected identity plasticity, which could be harnessed for regenerative purposes.

Regenerated RGCs connect retina to brain by long-distance projection of axons into various brain visual areas, even in animals where original RGCs and axons have been damaged. These reveal that the adult mammalian visual system remains a remarkable ability of reconnecting neural circuits. This opens opportunities for vision restoration by regenerating lost neurons and rebuilding the visual circuitry. Given that RGCs represent only a small population (about 1%) of retinal neurons, successful generation of functional RGCs by reprogramming other retinal neuronal lineages on site provides a promising therapeutic strategy for vision restoration in patients.

## ACKNOWLEDGEMENTS

The authors thank Xi Lin for technical help. This research is supported by NSFC (Grant # 81670880) to H. Liu.

## AUTHOR CONTRIBUTIONS

Conceptualization, H. L. and Y. Y.; Formal Analysis, X. W., Z. Z., and X. W.; Investigation, X. W., Z. Z., H. Z., X. W., W. Z., N. Q., Z. C., L. L., C. F., Z. Y., and X. L. Resources, H. L., H. Z., and X. W.; Writing - Original Draft, H. L.; Visualization, X. W., Z. Z., H. Z., X. W., and H. L.; Supervision, H. L. and Y. Y. Project Administration, H. L.; Funding Acquisition, H. L.

## STAR METHODS

### RESOURCE AVAILABILITY

#### Lead Contact

Further information and requests for resources and reagents should be directed to and will be fulfilled by the Lead Contact, Hongjun Liu.

#### Materials Availability

All unique resources generated in this study are available upon reasonable request to the lead contact with a completed Materials Transfer Agreement.

#### Data and Code Availability

This study did not generate/analyze unique datasets/code.

### EXPERIMENTAL MODEL AND SUBJECT DETAILS

#### Mice

The *Lgr5^EGFP-IRES-CreERT2^* knock-in mouse strain, the *Pvalb^CreERT2^* knock-in mouse strain, and the *Rosa26-tdTomato* reporter mouse strain were obtained from the Jackson laboratory. *Lgr5^EGFP-IRES-CreERT2^* mice and *Pvalb^CreERT2^* mice were crossed with *Rosa26-tdTomato* mice to generate *Lgr5^EGFP-IRES-CreERT2^; Rosa26-tdTomato* mice and *Pvalb^CreERT2^; Rosa26-tdTomato* mice, respectively.

The *Prokr2^CreERT2^* mouse strain was generated by homologous recombination using the CRISPR/Cas9 technology. Briefly, Cas9 mRNA, sgRNA and a donor vector plasmid were mixed and injected into the pronucleus of fertilized eggs from C57BL/6J mice. The donor vector plasmid was designed to insert the coding region of CreERT2 followed by a PolyA sequence into the ATG start codon of the Prokr2 locus. The injected zygotes were cultured until blastocyst stage by 3.5 days, and were subsequently transferred into uterus of pseudopregnant females. F0 mice with correct genome targeting were further crossed with C57BL/6J mice to generate F1 *Prokr2^CreERT2^* mice. *Prokr2^CreERT2^* mice were crossed with *Rosa26-tdTomato* mice to generate the *Prokr2^CreERT2^; Rosa26-tdTomato* mice.

All mice were housed in an animal facility with a 12-hour light/12-hour dark cycle. Animal experiments were conducted in both male and female mice of 8-12 months of age, and all animal experiment procedures were approved by the Animal Care and Use Committee at ShanghaiTech University.

#### Cell line

The HEK293 cell line used for AAV production was obtained from the Cell Bank of Shanghai Institute of Biochemistry and Cell Biology (SIBCB), Chinese Academy of Sciences, and cultured in DMEM with 10% FBS in the presence of penicillin/streptomycin in a 37 ° C incubator with 5% CO_2_.

### METHOD DETAILS

#### Construction and production of AAV vectors

Coding sequences of mouse Atoh7, Brn3B, Sox4, Sox11, Ils1 and EGFP were sub-cloned into the CAG-driven Cre-dependent expression vector (Addgene #22222), replacing the original Arch-GFP sequence. To co-express a transcription factor and EGFP from a single AAV vector, a P2A fragment was placed between the two coding sequences.

For AAV viral particle production, HEK293T cells were transfected with the AAV transgene plasmid, pAAV7m8 serotype plasmid and the pHelper plasmid using PEI. Cells were collected 48-72 hours later. Viral particles were purified with Iodixanol density gradient centrifugation, and tittered by qPCR.

#### Intravitreal AAV injection

Mice were anesthetized by IP injection of a mixture of ketamine (80 mg/kg) and xylazine (8 mg/kg), and their pupils were dilated with a topical administration of Phenylepherine Hydrochloride ophthalmic solution (2.5%). After a brief topical anesthesia with 0.5% Proparacaine Hydrochloride eye drop, a cornea puncture was made to reduce intraocular pressure, and a 1.5 ul of AAV viral particles was injected into the vitreous space with a 34-gauge needle. For injections of AAV mixtures, each AAV was first diluted to 1×10^12^ particles/ml before mixing.

#### Glaucoma model

Mouse RGCs were damaged using an intraocular pressure increase (IPI)-induced ischemia/reperfusion (I/R) model that mimics acute angle closure glaucoma in clinic. With minor modifications of a previously reported protocol (Hartsock et al., 2016), the ocular anterior chamber of mice was annulated with a needle, which is connected through a tube to an elevated saline (with 0.1% Heparin) reservoir. By elevating the height of the saline reservoir to 150 cm above the eye, the inner retinal blood flow was halted (ischemia). The needle was removed to install the circulation (reperfusion) 60 minutes later. This protocol causes degeneration of all RGC axons and death of other retinal neurons. To prevent other retinal neurons from apoptosis, a solution of Rock inhibitor Ripasudil hydrochloride dehydrate (0.4% in PBS) was administrated to the eye surface of mice once a day.

#### Immunohistochemistry and imaging

After being transcardially perfused with saline (0.9% NaCl in ddH2O) and subsequently 4% PFA, eyes, optic nerves and brains of mice were collected and post-fixed in 4% PFA for 24 hours. Eyes and brain tissues were placed in 30% sucrose for cyroprotection, and sectioned using a Microtome Cryostat at thickness of 10 and 30 μm, respectively. Immuno-histochemical stainings were performed according to a standard protocol. The following antibodies were used: rabbit-anti-RBPMS (Abcam, 1:400) to label RGCs, mouse-anti-Brn3a (Santa Cruz Biotechnology, 1:200) to label RGCs, rabbit anti-SMI-32 (Abcam, 1:400) to label α-RGCs, rabbit anti-melanopsin (Abcam, 1:500) to label ipRGC, rabbit anti-CART (cocaine- and amphetamine-regulated transcript) (Phoenix Peptide,1:2500) to label ON-OFF DSGCs, mouse anti-PSD95 (Abcam, 1:400) to label postsynaptic cell membrane. For secondary detections, Alexa Fluor 647 donkey anti-rabbit (IFKine™,1:400), Alexa Fluor 647 donkey anti-mouse (IFKine™,1:400), or Alexa Fluor 488 donkey anti-rabbit (Abcam, 1:400) were used. Immuno-stained tissue sections were imaged with a Zeiss LSM880 confocal microscope, a Nikon spinning disk (CSU W1 Sora) confocal microscope or a STED SP8 microscope.

#### *In vivo* calcium imaging

For surgery, mice were anaesthetized with urethane (1.5 g/kg), and placed in a stereotaxic device with eyes covered with ophthalmic ointment. A custom titanium head-plate was bonded to the skull with black dental cement (Fe3O4 was added to block light), roughly centered on lambda, parallel to the long axis of the mouse. A 3-mm craniotomy was performed over the posteromedial SC and inferior colliculus, and a coverslip with 3 mm diameter was then gently pressed upon the dura and the craniotomy was sealed with black dental cement. A piece of black-out cloth was attached on the head-plate to avoid light contamination by the visual stimulation during functional two-photon imaging.

Visual stimuli were generated using the Matlab (Mathworks) function Psychtoolbox and displayed on a corrected 17’ LCD monitor (Dell, 1280 by 1024 pixels, 75 Hz refresh rate) positioned 15 cm from the contralateral eye. The stimuli were a full screen of sine-wave drifting gratings presented on a gray homogeneous background (spatial frequency: 0.05 cycles/°, temporal frequency: 2 Hz). The gratings were presented for 5 repeats with 1s duration and 1-2 s interstimulus interval. The stimuli were drifted in 8 directions orthogonally to 4 orientations at regular intervals of 45°.

Two-photon imaging of fluorescence from axonal terminals was monitored with a customized LotosScan microscope (LotosScan, Suzhou Institute of Biomedical Engineering and Technology) and coupled with a mode-locked Ti:Sa laser (Chameleon VISION-S, Coherent). The excitation wavelength was fixed at 920 nm. Imaging was performed using a 40X, 0.8 NA objective (Nikon). The beam size was large enough to overfill the back aperture of the 40X objective. Images were acquired at a frame rate of 50 Hz (480 x 240 pixels, 0.225 μm/pixel).

Images were analyzed in Matlab (Mathworks) and ImageJ (National Institutes of Health). For correcting lateral motion in the imaging data, a rigid-body transformation based frame-by-frame alignment was applied by using Turboreg software (ImageJ plugin). Terminals were identified by hand on the basis of size, shape, and brightness. Individual terminal time courses were extracted by averaging pixel intensity values within terminal masks in each frame. If brain pulsation were evident during imaging, these data were not used. Neuropil signal was subtracted by using the method previously reported (Kerlin et al., 2010). After this correction, responses (Ft) to each stimulus presentation were normalized by response in the 0.2s immediately before the stimulus onset (F0). For each stimulus, the mean change in fluorescence (ΔF/F) was calculated by averaging responses to all stimulus conditions and trials. Visually responsive cells were defined by ANOVA across blank and stimulus presentation periods (P<0.05).

#### *In vitro* whole-cell patch clamp recording

Deeply anesthetized mice were transcardially perfused with an ice-cold oxygenated (95% O_2_, 5% CO_2_) cutting solution containing 92 mM Choline-chloride, 2.5 mM KCl, 1.2 mM NaH2PO4, 30 mM NaHCO3, 10 mM MgSO4, 0.5 mM CaCl2, 25 mM Glucose, 5 mM Na-ascorbate, 3 mM Sodium Pyruvate and 2 mM Thiourea. The pH value of the cutting solution was adjusted to 7.3-7.4 by adding concentrated HCl and the osmolarity was adjusted to 310-315 mOsm. After being removed from the skull, brain tissues containing the SC region were cut into 300 μm coronal slices within the cutting solution, using a vibrating blade microtome (VT1200 S, Leica Biosystems). Slices were then incubated in the same cutting solution at 31-32 °C for 15 minutes, before being transferred into a holding chamber containing room-temperature oxygenated holding solution (92 mM NaCl, 30 mM NaHCO3, 1.25 mM NaH2PO4, 2.5 mM KCl, 2 mM MgSO4, 2 mM CaCl2 and 25 mM Glucose, 20 mM HEPES, 5 mM Na-ascorbate, 3 mM Sodium Pyruvate, and 2 mM Thiourea, with a pH value of 7.3-7.4 and a osmolarity value of 310-315 mOsm). After storing for one hour, the slices were transferred into a recording chamber containing room-temperature oxygenated recording solution (119 mM NaCl, 24 mM NaHCO3, 1.25 mM NaH2PO4, 2.5 mM KCl, 2 mM MgSO4, 2 mM CaCl2 and 12.5 mM Glucose). Three to five slices containing the SC region were typically produced from one animal. Recordings were taken from brain slices containing the middle SC region.

Whole-cell patch clamp recordings of synaptic responses were made using a 2-4 MΩ glass pipettes with an internal solution of 125 mM K-gluconate,20 mM KCl, 0.5 mM EGTA, 10 mM HEPES-NaOH, 10 mM P-Creatine, 4 mM ATP-Mg, and 0.3 mM GTP (pH 7.3). Blue stimulation light was produced by a 470 nm LED (Thorlabs, 35mW/mm^2^) and applied through an 40X objective (OLYMPUS). Stimulation duration at 5 ms was found to be able to saturate postsynaptic responses recorded. Neurons had input resistances in a range of 1-5 GΩ and series resistances less than 20 MΩ. Recordings were performed with the following protocol: The membrane potential was first held at −70 mV to record the light-evoked AMPA receptor-mediated synaptic currents (NMDA receptors were presumably blocked by magnesium at this holding potential). The membrane holding potential was then switched to +55 mV to record a mixture of AMPA and NMDA receptors-mediated currents. Under this condition, AMPA receptor antagonist CNQX (10 mM) was then added to the recording solution to block AMPA receptor-mediated synaptic currents, allowing detection of NMDA receptor-mediated EPSCs. Next, the recording was switched to current clamp mode to detect action potential. Applications of the AMPA receptor antagonist CNQX (Tocris) and the NMDA receptor antagonist D-APV (Tocris) were performed by adding respective drugs into the bathing recording solution. All recordings were made with an Axon700B amplifier and digitized using a Digidata1440 analog-to-digital board. Stimulation and data acquisition were performed with the pClamp software and digitized at 50 kHz. All equipment and software are from Axon Instruments/Molecular Devices (Molecular Devices, CA).

### QUANTIFICATION AND STATISTICAL ANALYSIS

The number of regenerated RGCs was quantified by counting tdTomato^+^ axons in optic nerves of the *Lgr5^EGFP-IRES-CreERT2^; Rosa26-tdTomato* mice and EGFP^+^ axons in optic nerves of the *Prokr2^CreERT2^; Rosa26-tdTomato* mice, six weeks after viral injection. Eight optic nerves from four male mice in each group were counted. Data with statistical analysis were presented as mean ± standard errors. Differences between two groups were compared using a two-tailed Student’s *t*-test. To quantify alterations of brightness of regenerated RGC terminals in the SC region, terminals were hand-picked based on the size, shape, and brightness. Individual terminal time courses were extracted by averaging pixel intensity values within terminal masks in each frame. After subtracting neuropil signal, responses (Ft) to each stimulus presentation were normalized by response in the 0.2s immediately before the stimulus onset (F0). For each stimulus, the mean change in fluorescence (ΔF/F) was calculated by averaging responses to all stimulus conditions and trials. Visually responsive cells were defined by ANOVA across blank and stimulus presentation periods (P<0.05).

